# Contact-dependent cell-cell communications drive morphological invariance during ascidian embryogenesis

**DOI:** 10.1101/238741

**Authors:** Léo Guignard, Ulla-Maj Fiuza, Bruno Leggio, Emmanuel Faure, Julien Laussu, Lars Hufnagel, Grégoire Malandain, Christophe Godin, Patrick Lemaire

**Affiliations:** CRBM, Université de Montpellier, France; Inria project-team Virtual Plants, CIRAD, INRA, Université de Montpellier, France; Janelia Research Campus, Howard Hughes Medical Institute, 19700 Helix drive, Ashburn, VA, USA; European Molecular Biology Laboratory, Cell Biology and Biophysics Unit, Meyerhofstrasse 1, 69117, Heidelberg, Germany; Institut de Biologie Computationnelle, IBC, Université de Montpellier, France; IRIT, CNRS, INPT, ENSEEIHT, Universités de Toulouse I et III, France; Université Côte d’Azur, Inria, CNRS, I3S, France

## Abstract

Canalization of developmental processes ensures the reproducibility and robustness of embryogenesis within each species. In its extreme form, found in ascidians, early embryonic cell lineages are invariant between embryos within and between species, despite rapid genomic divergence. To resolve this paradox, we used live light-sheet imaging to quantify individual cell behaviors in digitalized embryos and explore the forces that canalize their development. This quantitative approach revealed that individual cell geometries and cell contacts are strongly constrained, and that these constraints are tightly linked to the control of fate specification by local cell inductions. While in vertebrates ligand concentration usually controls cell inductions, we found that this role is fulfilled in ascidians by the area of contacts between signaling and responding cells. We propose that the duality between geometric and genetic control of inductions contributes to the counterintuitive inverse correlation between geometric and genetic variability during embryogenesis.

## INTRODUCTION

Within each animal species, embryonic development is highly reproducible, ensuring the production of a complex organism with precisely arranged and shaped organs and tissues. This constancy of embryogenesis against genetic polymorphism and fluctuating environmental conditions is critical for the perpetuation of the species, and has been referred to as developmental canalization (*1*).

Although tissue-scale embryonic reproducibility results from the careful orchestration of cell behaviors during development (*2*), it does not imply that individual cells behave reproducibly from embryo to embryo. Rather, tissue-level invariance is in most species an emergent property, which results from the averaging of the variable (*3*) or even stochastic (*4*) behaviors or mechanical properties (*5*, *6*) of individual cells.

Some invertebrate species, including most nematodes (*7*) and ascidians (*8*), exemplify an extreme form of canalization. They develop in such a highly stereotyped manner that the position and fate of individual embryonic cells, as well as the orientation and timing of their divisions, show very little variability between individuals, leading to essentially invariant embryonic cell lineages (*9*, *10*). This cellular reproducibility of wild type development is robust to the unusually high level of genetic polymorphism found in nematodes and ascidians (*11*–*13*). Early ascidian cell lineages and embryo geometries have even remained essentially unchanged since the emergence of the group, around 500 million years ago, despite extensive genomic divergence (*8*, *14*). Because of the stereotypy of their early development and remarkable ability to buffer genetic divergence, ascidians constitute an attractive system to understand the mechanistic origin of extreme canalization.

Canalization has mostly been analyzed through the prism of the developmental Gene Regulatory Networks (GRNs) that drive and coordinate development. The current view is that reducing gene expression variability is a key feature of canalization achieved through the use of specific GRN architectures (*15*–*18*) or through the folding or stabilization of signal transduction proteins by specific chaperones (*19*). Gene regulatory networks are, however, much less evolutionary robust than morphologies in both ascidians (*14*) and nematodes (*20*). Additionally, in the ascidian ***Ciona intestinalis***, the majority of genes show variable maternal expression between individuals (*21*). As extreme canalization of embryonic morphologies is observed despite variable gene expression and gene regulatory network, it is unlikely to be explained solely by the canalization of regulatory gene expression.

What else could explain cell-scale geometric invariance? The larvae of most species with invariant cell lineages are made of only a few hundreds to thousands of cells. This reduced complexity could favor stereotypy, but it is not ***per se*** sufficient to explain it: the marine nematode, ***E. brevis***, develops with similar cell numbers as ***C. elegans***, but with variable, indeterminate cell lineages (*8*). Stereotypy does also not result from the use of qualitatively different developmental strategies from those of other animals: while the precise partitioning during cell division of autonomously-acting localized maternal development was initially thought to drive stereotyped development (*23*, *24*), cell communication is now recognized as equally important in embryos with invariant or variable cell lineages (*25*).

In this study, we test the hypothesis that the combination of short-range cell communication events involved in ascidian fate specification and reduced cell numbers, strongly constrains embryonic geometries. For this, we first developed experimental and algorithmic methods to perform long-term multi-view live imaging of developing ascidians and to automatically extract the precise geometries of cells and cell-cell contacts with unsurpassed accuracy and used these to estimate their degree of variability between embryos. We then combined cell contact geometries with an atlas of signaling gene expression to build a computational model of cell inductions, whose robustness to geometric and genetic variations we studied. Our results reveal that cell fate specification by surface-dependent inductions imposes strong local constraints on the areas of contact between communicating cells. This strongly canalizes embryonic geometries, while putting lesser constraints on the evolution of the genome.

### A high-resolution geometric atlas of embryonic cell shapes and interactions

Using confocal multiview lightsheet microscopy (*26*), we imaged live ***Phallusia mammillata*** embryos with fluorescently-labeled plasma membranes every two minutes from 4 orthogonal viewing directions and for several hours, without compromising their development. High quality 4D datasets with isotropic spatial resolution were obtained after fusion of the four 3D images captured at each time point (Figures S1, S2). They extend from the 64-cell stage to the initial tailbud (Embryo1) and late neurula (Embryo2) stages respectively, covering two major morphogenetic processes, gastrulation and neurulation (Supp. datasets, Video S1, Figure 1), through up to 4 cell divisions.

**Figure 1:**
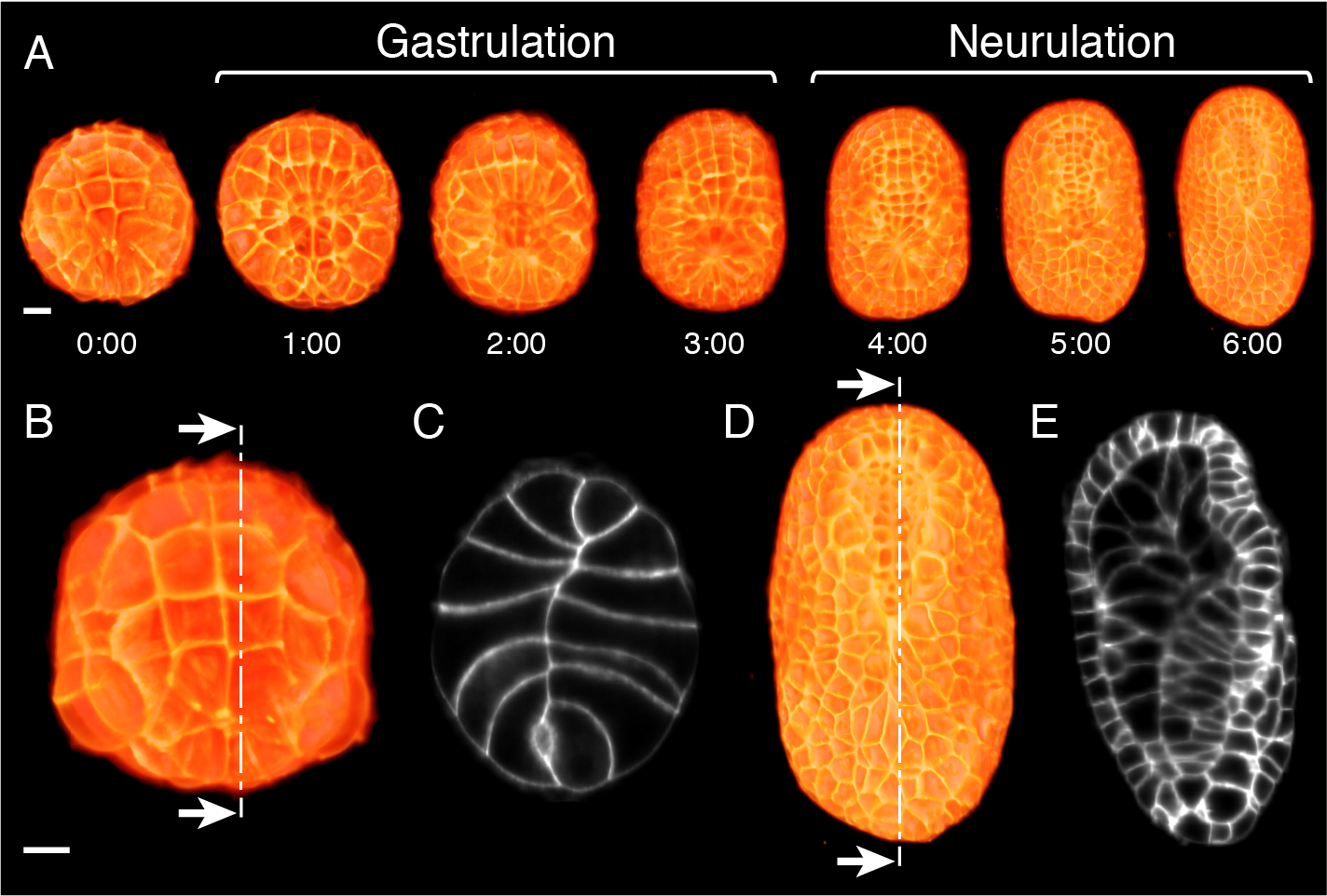
High-resolution multiview lightsheet live imaging of *Phallusia mammillata* embryos. A) Vegetal/dorsal views of 3D rendering at the indicated time points of the imaged embryo after fusion of the images taken along the 4 angles of views. B) Vegetal view of the embryo at the 64-cell stage (t = 1). C) Sagittal section along the plane shown on B. D) Dorsal view of the embryo at time t = 180. E) Sagittal section along the plane shown on D. A-E, anterior is to the top. Scale bar: 20μm.

Systematic segmentation and long-term tracking of all membrane-labeled cells of a developing embryo is notoriously challenging *(*27*)*. Building on our previous low-throughput MARS-ALT pipeline (*28*)(Figure S3-6), we developed a novel algorithm, ASTEC, for Adaptive Segmentation and Tracking of Embryonic Cells (Figure 2A), able to faithfully segment whole cells and track cell lineages over long periods of time with high temporal resolution. ASTEC is a single pass algorithm, which simultaneously segments and tracks cell shapes by propagating information over time, a strategy pioneered for nuclear labels by Amat and colleagues (*29*). ASTEC is initiated with a manually curated segmentation of the first time point, and iteratively projects segmentations from one time point to the next (Figures S7-11 and Supp. information), then detects cell division events between consecutive time points. It produces a segmentation of all embryonic cells present at each time point, and for each cell “snapshot” (the image of a cell at a given time-point) the identity of its progeny at the next time point, from which global cell lineages and temporal dynamics of individual cell shapes and physical cell-cell contacts can be reconstructed.

**Figure 2:**
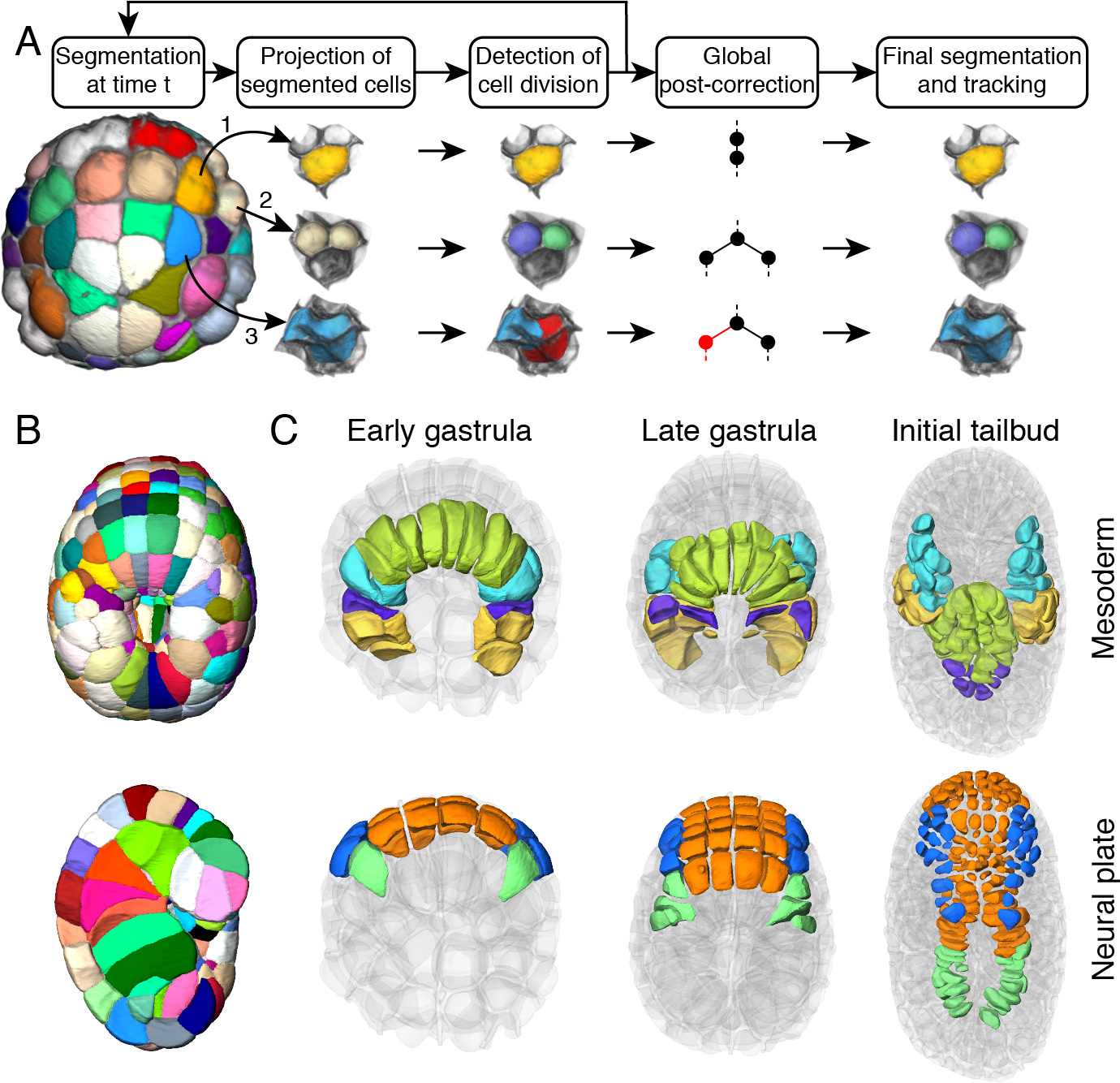
ASTEC segmentation of *Phallusia mammillata embryos*. A) The ASTEC pipeline. Three different cases are illustrated: a non-dividing cell (yellow), a dividing cell (white), an oversegmented non-dividing cell (blue) corrected during post-correction. For more details see Figure S7. B) Segmentation of ASTEC1 at t=76. Top: view from the dorsal side. Bottom: sagittal section. Colors are arbitrary. C) Evolution of the position of mesoderm and neural progenitors in ASTEC1 at the early gastrula (t=35), late gastrula (t=80) and initial tailbud (t=170) stages. Mesoderm: yellow, B-line mesenchyme; purple, secondary notochord; green, primary notochord; cyan, Trunk Lateral Cells; muscle is not represented. Neural plate: green, lateral tail nerve cord; blue, dorsal anterior neural plate; orange, ventral anterior neural plate. Anterior is to the top. For more details see Figure S13.

The resulting segmentation (Figure 2B, Supp. Dataset, Video S2) and global lineage trees of Embryo1, ASTEC1, contain a total of 58454 digital 3D cell “snapshots”, describing the behavior in time of 1342 individual cells generated by 639 cell division events. Integrated with the known fate of early blastomeres *(*10*)*, this geometric description of a developmental program allows to track with high temporal and spatial resolution the position, geometry, contacts and lineage history of every embryonic cell (Figures 2C, S12, 13, Video S3). Quality assessment of the segmentation and lineage of ASTEC1 indicated that 99% of voxels were assigned to the right cell (Figure S14), that cell volumes were consistent between matching cells in the left and right halves of the embryo and that the sum of the volumes of daughter cells closely corresponded to the volume of their mother (Figure S15). The pattern of rounding up of cells as they approached mitosis indicated that cell divisions were detected with a temporal accuracy of 2 minutes (Figure S16, S17). No programmed cell death was detected, as described in other ascidian species *(*30*)*. The segmentation and tracking accuracy of the second embryo, ASTEC2, obtained by running ASTEC with the same parameters, was comparable (see Supp. information). The ASTEC1 and ASTEC2 datasets (Supp. Data sets) constitute to our knowledge the first systematic quantitative descriptions of the dynamics development including cell lineages, and the geometry and interactions of whole cells, rather than cell nuclei, across a large fraction of a metazoan developmental program.

### Stereotypy of ascidian development

Although ascidians are considered a textbook example of invariant development, few studies have attempted to go beyond qualitative observations to quantify inter-embryo variability. One such study indicates that at least the notochord shows evidence of stochastic cell intercalation during the tailbud stages (*31*). Our geometric atlas gave us the opportunity to quantitatively assess the variability of ascidian cell lineages, cell geometries and cell neighborhoods during gastrulation and neurulation.

We first compared the temporal progression in cell numbers in our two ASTEC-reconstructed embryos and in an independently imaged, manually-curated, ***Phallusia mammillata*** embryo whose nuclei rather than membranes were fluorescently labeled (BioEmergences embryo) (*32*). Cell numbers were remarkably consistent between embryos until the late neurula stage (7 hpf at 18°C), with only slight differences beyond (Figure 3A). Consistently, the structure of cell lineage trees originating from matching cells of early gastrula progenitors of ASTEC1 and the BioEmergences embryos, or from equivalent left and right cells within each embryo, differed by less than 10% when compared with a tree edit metric (*33*) (Figures 3B and S18-19). Lifespans of matching cells also only differed marginally, by less than 10% on average, between embryos or between the right and left halves of each embryo (Figures 3C, S21).

**Figure 3:**
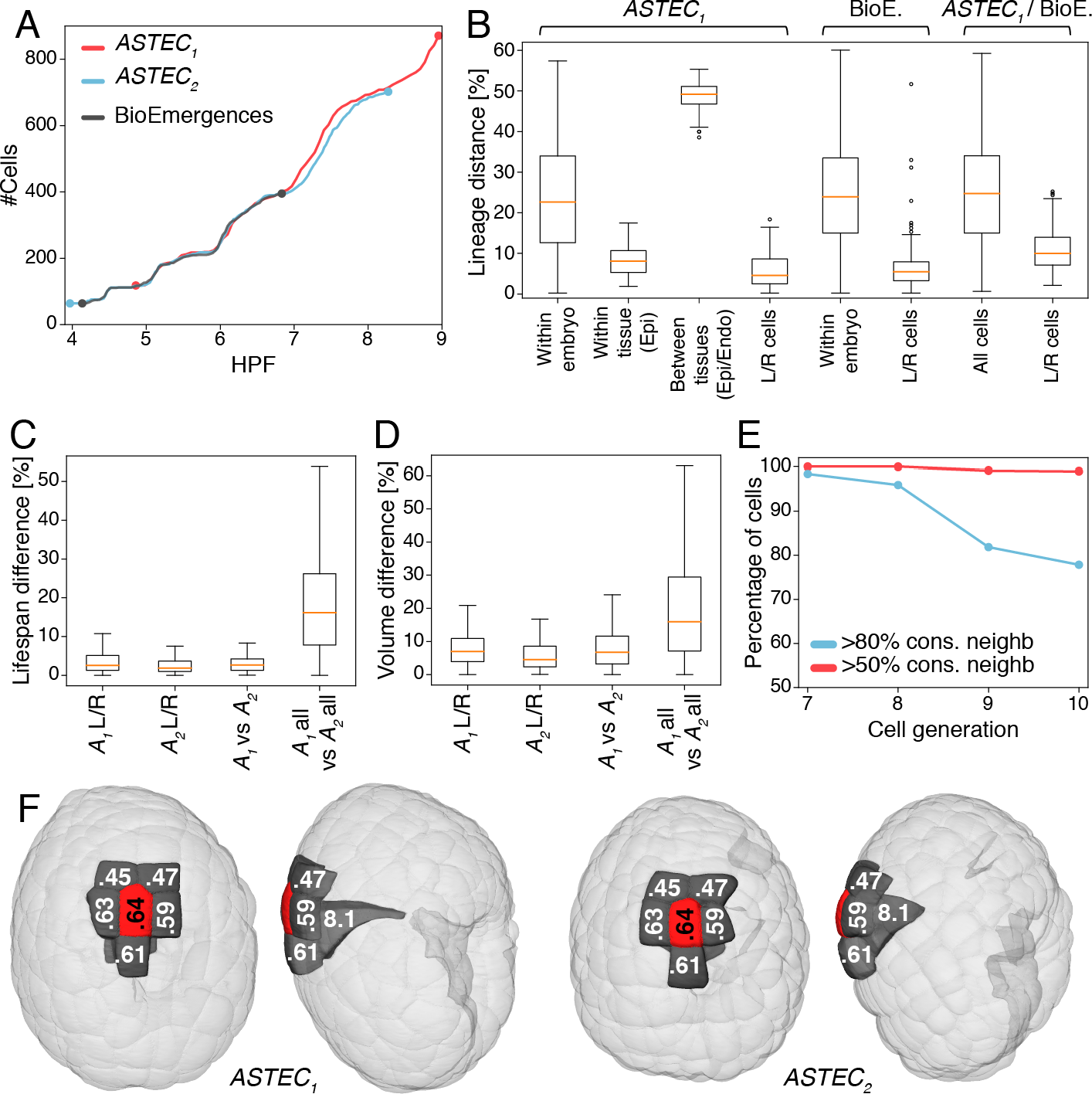
Stereotypy of ascidian development. A) Comparison of the evolution in time of cell numbers between three individual embryos after linear temporal rescaling (see Supp. data). B) Distributions of pairwise lineage distances between trees originated at the 112-cell stage from the ASTEC1 and BioEmergences embryos. Within Embryo, distribution of all pairwise distances; Within tissue (epi), pairwise distances between epidermis and endoderm; (Between tissues), pairwise distances between trees from matching bilateral cells (L/R cells) and for both ASTEC and BioEmergences lineage trees (All cells). For more details, see Figure S19. Boxes show the first, second and third quartiles, whiskers the range up to 1.5 interquartile. C-D) Distributions of the relative differences in lifespans (C) and Volumes (D) between matching cells within and between ASTEC1 and ASTEC2 embryos. See also figs S20 and S21. E) Percentage of cells with 80% or 50% of common physical neighbors (contact >5% of the cell surface) between matching cells in ASTEC1 and ASTEC2 embryos. See also Figure S23. F) Example of a cell with perfectly conserved neighborhood in the two ASTEC embryos. Note the conservation of the spatial position of the cell in both embryos. Left: ventral view, right: lateral view. Light grey cells are translucent.

Temporal invariance of cell lineages does not necessarily imply geometric invariance (*34*). We therefore next assessed geometric variability in ***Phallusia***. While the distribution of cell volumes at a given time point was broad, the volume difference of matching cells was inferior to 10% on average, both between ASTEC1 and 2 and between the bilateral halves of each embryo (Figures 3D and S20). Cell organization was also shared. More than 80% of matching cells in ASTEC1 and ASTEC2 shared at least 80% of their physical neighbors (Figure 3E). Most cell-cell contacts shared between embryos persisted for the whole life of the cells and most cells kept the same neighbors throughout their life (Figure S22). As a consequence, we found no evidence of individual cell migration or cell intercalation up to the initial tailbud stage. This topological invariance extended to the invariance of the measure of areas of contacts between cells. In 80% of matching cells, the average variation of surface of contact to shared neighbors was smaller than 20%, again both between bilateral halves of each ASTEC embryo, and between embryos (Figures 3E, S23). This conservation of local neighborhoods translated into a conservation of the global spatial position of each cell with respect to embryonic axes (Figure 3F).

Thus, cell divisions, cell geometries, cell arrangements and positions are highly reproducible between individual ***Phallusia*** embryos until at least the late neurula stages. The comparable extent of inter-embryonic and bilateral intra-embryonic variability suggests that developmental noise, rather than genetic variation, is an important driver of this variability.

### Asymmetric cell divisions couple fate specification to embryonic cell geometries

To identify the causes of ***Phallusia*** developmental invariance, we first considered the cellular basis of embryonic morphogenesis. Embryonic morphogenesis is built from few elementary cell behaviors (*2*). In the absence of cell migration/intercalation, cell growth and cell death, ***Phallusia*** morphogenesis must be primarily driven by oriented cell division, geometrically unequal divisions and cell shape changes. Asymmetric cell divisions, defined as divisions generating differently fated daughter cells, can couple fate specification to morphogenesis as some asymmetric divisions also control the relative position in space (oriented divisions) or volume (unequal cleavages) of daughter cells (*35*). Using our geometric atlas, we set out to systematically identify asymmetric cell divisions and assessed their geometric consequences.

We first scanned cell lineages for a signature of asymmetric cell divisions. Using our tree edit distance, we hierarchically clustered all 64-cell ***Phallusia*** progenitors in ASTEC1 on the basis of the similarity of the cell lineage trees they seeded (Figures S24, S23). The majority of cells with identical or similar tissue fates clustered together, indicating that the mitotic history of ascidian embryonic cells is strongly correlated to their larval fates. We therefore reasoned that a comparison of the structure of the sublineage trees originating from two sister cells may identify sisters with distinct fates, and thus originating from candidate asymmetric cell divisions (Figure 4A). Figure 4B shows the distribution of distances between the cell lineage trees originating from 108 bilateral pairs of sister cells between the 64-cell and mid-gastrula stages (time point 124 for ASTEC1). Only 35/108 divisions gave rise to lineages trees differing by more than 10%, a subset including 19/22 known fate restriction events (Table S1 and Figure S26). This approach also identified 16 novel asymmetric cell divisions. 14 of these divisions were found in tissues known to give rise to several larval or juvenile mesodermal tissues, such as the mesenchyme and trunk lateral cells (TLC) (*36*–*38*), or to show strong cellular heterogeneity, such as the central nervous system (*39*, *40*) or the tail epidermis (*41*). We thus used lineage tree structure asymmetry as a marker for asymmetric cells divisions.

**Figure 4:**
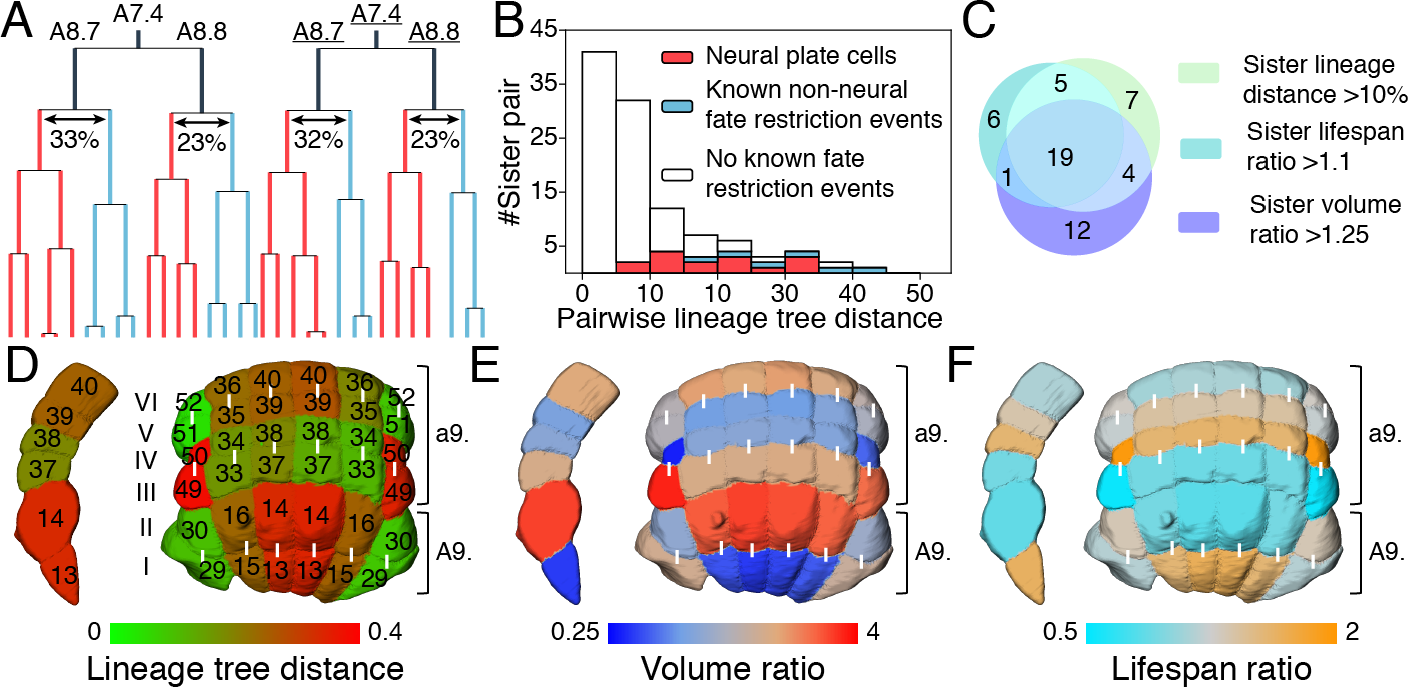
Relationships between fate restriction, unequal cleavage and unequal life spans. A) Example of the asymmetric division of the A8.7 and A8.8 posterior neural progenitors. The trees seeded by the two daughters of A8.7 and A8.8 are colored differently. Percentage values indicate the tree-edit distance between the lineages seeded from the daughters of A8.7 and A8.8 cells. Note the similarity of the lineages originating from the left and right A7.4 progenitors. B) Distribution of the cell lineage distance between sister cells born between time points t= 1 and t=124. C) Venn diagram showing the overlap of cell divisions leading to high sister lineage distance, sister lifespan ratio and sister volume ratio for cells generated between time points t= 1 and t= 124. D, E, F) Dorsal and lateral view of the a- and A-line derived neural plate color-coded for lineage tree distance (D), Volume ratios (E) and lifespan ratios (F). White bars link sister cells. The identity of 9th generation cells is indicated on panel E. Lineage tree distances are features of a cell pair, while ratios are features of individual cells.

We next explored the geometrical impact of asymmetric divisions (Figure 4C and Table S1). While unequal cleavages are infrequent in ***Phallusia*** (Figure S27A, B), 23/35 asymmetric divisions were also geometrically unequal. Interestingly, in geometrically unequal divisions, the smaller daughter generally had a longer lifespan (Figure S27C), in agreement with a general anti-correlation found between cell volumes and lifespans in our dataset (Figure S27D), and in ***C. elegans*** (*42*). Unequally dividing cells also significantly departed from the default Hertwig rule (*43*) for the orientation of the division (Figure S27E). Co-occurrence of asymmetric divisions, geometrically unequal cleavage and differential lifespans was particularly strong in the mid-gastrula neural plate (Figure 4D-F).

Thus, up to the mid-gastrula stage, around 30% of all cell divisions are asymmetric and the majority of these divisions couple the fate specification process to the geometry, relative position and lifespan of daughter cells.

### Geometric control of differential cell inductions

We next analyzed whether, conversely, cell geometry and neighborhood relationships could impact fate specification. In ascidians, most studied asymmetric cell divisions are triggered by contact-dependent cell communication events, which either polarize the mother cell or differentially induce its daughters (*44*). The frequency of asymmetric divisions we detected suggests that contact-dependent inductions may collectively impose a global constrain on embryonic geometries, thereby promoting stereotypy. To test this idea, we built computational models of differential cell induction integrating geometrical features extracted from the ASTEC1 dataset with the expression pattern of signaling genes, which in ascidians have been systematically determined with a cellular resolution up to the onset of gastrulation (*45*, *46*). Simulated cell inductions were then confronted to a ground truth of experimentally characterized differential induction events and uninduced cells (Table S2).

Extracellular ligands or antagonists for only six major signaling pathways (FGF, Ephrin, Wnt, Bmp, Nodal and Notch) show patterned expression during the cleavage and early gastrula stages (Figure S29). As receptors and intracellular components of these pathways are maternally expressed, we considered them ubiquitous. All cells were therefore considered competent to respond to all pathways, and ligand availability was the limiting factor controlling inductions. We computed ligand availability by integrating the pattern of expression of the ligand gene with that of its extracellular antagonists, the presence of an antagonist at a cell interface fully blocking the action of the ligand at this interface.

First, we tested whether direct physical contact of a competent cell to a cell emitting a ligand could be sufficient to ensure its induction, irrespective of the area of contact (Figure 5A). We defined the signaling state of a cell as the combination of free ligands a cell is exposed to and compared the signaling states of sister cells to experimentally determined fate specification decisions (Figure 5B,C and S35). Cells were found to respond on average to 5.5 ligands and the resulting 25 cell signaling states from the 64-cell to the early gastrula stages could only explain 4/14 known differential induction events. Thus, a simple induction model based on the topology of physical cell contacts is insufficient to account for known asymmetric cell divisions.

**Figure 5:**
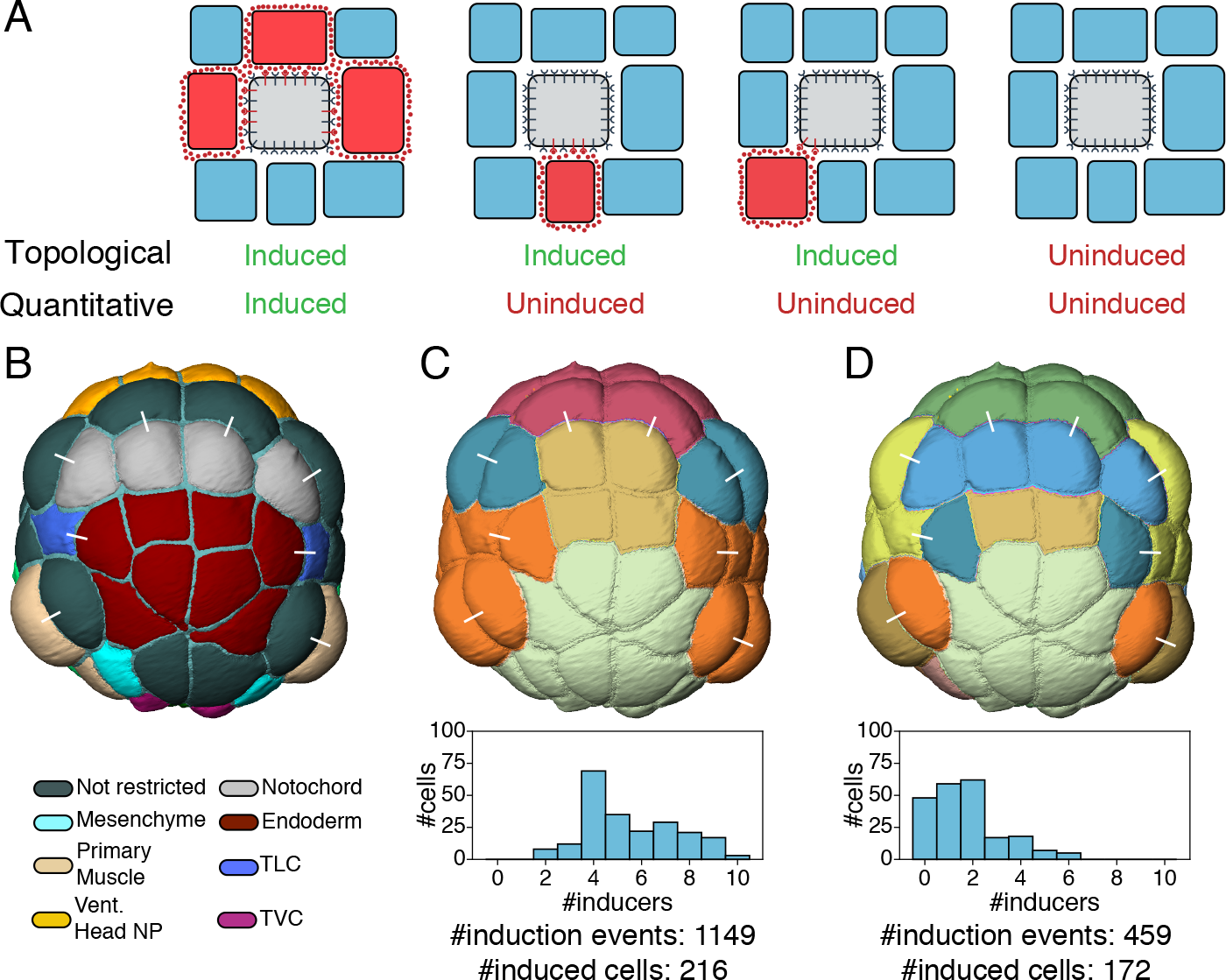
Models of cell-cell contact-dependent inductions. A) Two scenarios of contact-dependent cell inductions. In the “Topological cell contact” scenario, any responding cell (grey) contacting a ligand-emitting cell (red) is induced. In the “Quantitative cell contact areas” scenario, a cell is only induced if its area of contact with ligand-expressing cells is sufficiently large. B) Color-coded fate map of cells at the 64-cell stage. C) “Topological cell contact” scenario. Top: color-coded signaling states; Bottom: distribution of the number of pathways affecting a cell’s fate for cells born between time points 1 and 124. D) “Quantitative cell contact areas” scenario resulting from the computational model. Top: color-coded signaling states. Bottom: distribution of the number of pathways affecting a cell’s fate for cells born between time points 1 and 124. In B-D, a white bar links sibling cells. Colors in B, D are arbitrary. See also Figure S35 for an equivalent analysis at the 112-cell stage.

The analysis of FGF-mediated ascidian neural induction (*47*, *48*) and of Notch signaling during hair cell specification in the chick basilar papilla (*49*) suggest that cells can take into account the area of contact with ligand-expressing cells in the interpretation of inducing cues. The high spatial and temporal resolution of our digital embryo made it possible to computationally test the generality of such a hypothesis at the level of a whole embryo. For this, we developed a quantitative continuous model based on the law of mass action (Figures S28, S30). Through a set of coupled differential equations modeling ligand-receptor-effector cascade dynamics, this model computes for each embryonic cell, at each stage, and for each pathway, the fraction of activated intracellular effector as a function of the surface of the cell, or of the part of its mother it inherited, exposed to a ligand from this pathway. Under the hypotheses of the model (Supp. Information), the fraction of activated intracellular effectors computed for each pathway was a quasi-linearly increasing function of the area of exposure to free ligands (Figure S31).

To identify relevant induction thresholds, we designed a set of logical consistency rules (Figures S32, 33) transforming this continuous information into a binary map of differentially induced sister cells (Figure S28). We then explored the parameter space (Figure S34) to maximize the fit of the model with a set of experimentally characterized differential inductions up to the early gastrula stage (Table S2). The best sets of parameters identified (see Supp. Information and Figure S34) were consistent with published knowledge (e.g. induction time of 8 minutes for all pathways, similar as in (*50*) for FGF). Compared to the topological cell contact model, this quantitative model led to a reduction of the number of signaling pathways each cell was responding to (1.7 on average), while increasing the number of signaling states (38 between the 64 and early gastrula stages), which were highly correlated to differential inductions events (Figures 5D, S35). The model correctly predicted inducers for all known induction events while over-predicting differential inductions for only 12 out of 56 (21%) symmetric divisions (Figures 6A, S36 and Table S3). The fractions of activated intracellular effectors for each pathway, each sister cell pair and each stage are given in Figures S37-54.

This remarkable result was strongly dependent on the hypotheses and initial conditions of the model. The performance was, as expected, much degraded when we mimicked the topological contact model by uncoupling the number of activated receptors from the area of cell-cell contacts or upon randomization of the patterns of expression of ligand and inhibitor genes (Figures 6A, S55, 56 and Table S3). The model was also able to recapitulate experimental perturbations which had not been used in its training. Virtual inhibition of Ephrin signals in the model phenocopied the biological knock-down (*51*–*53*) (Figure 6A, S57 and Table S3). Finally, the results of the model were robust to naturally occurring evolutionary changes in embryo geometry. While egg diameters are uniform within each species, they range across ascidian species from about 100 μm to nearly 1 mm. Consistent with the idea that most molecular mechanisms are evolutionarily conserved, even between distantly-related ascidian species (*54*), predictions of the model were robust to a 4-fold uniform scaling of the surfaces of cell-cell contact (Figures S58, S59).

**Figure 6:**
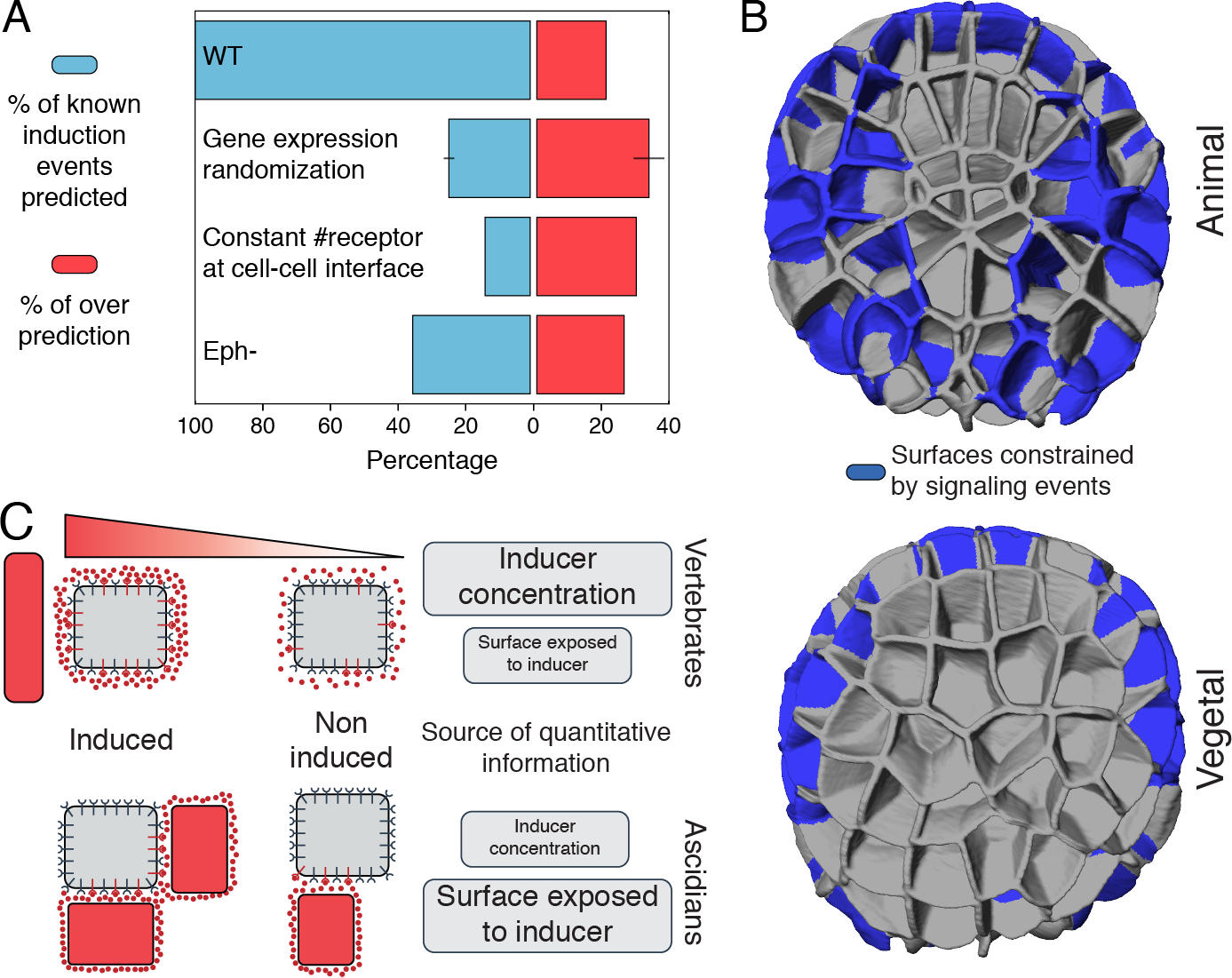
Predictions of the quantitative cell contact areas model. A) Percentage of known (blue) or overpredicted (red) cell induction events predicted by the model in Wild type (WT), and in situations of ligand expression randomization, constant available receptor situation and Eph receptor mutation. B) Visualization of the 112-cell stage cell-cell interfaces constrained by differential cell inductions. See also Figure S60 for equivalent data at the 64-cell stage. C) Conceptual models of vertebrate and ascidian embryonic cell inductions.

Taken together, our model of surface-dependent differential cell inductions, despite its simplicity, accounts with remarkable explanatory ability for early ascidian fate specification events up to gastrulation.

### Higher robustness to genetic than geometric perturbations

Our original aim was to shed light on the apparent paradox of a highly canalized and evolutionary conserved embryo morphogenesis in a context of rapidly divergent genomes, gene regulatory networks and gene expressions. We reasoned that if cells operate close to induction threshold values, small changes in embryo geometry might alter the induction or polarization status of some cells, thereby constraining a significant fraction of cell-cell interfaces. Indeed 32% of cells saw their polarization or induction status for at least one pathway change upon a +/−20% change in the surface of contacts they established with signaling cells. This local geometric sensitivity of inductions imposes a global constraint on cell interfaces (Figures 6B and S60), indicating that surface-dependent cell inductions have a strong canalizing effect on embryo geometries. By contrast, similar changes in the magnitude of ligand gene expression had a much smaller effect on cell inductions and polarizations (Figure S61).

The reliance of ascidian embryonic fate specification on the areas of contacts with signaling cells therefore ensures both high canalization of morphogenesis and more relaxed constraints on genomic information.

## Concluding remarks

Systematically and quantitatively tracking the temporal dynamics of multiple cellular properties, including shape, and cell interactions in all cells of a developing organism has been a dream for generations of developmental biologists. Previous dynamic atlases of development with cellular resolution have so far mostly been generated from embryos with fluorescently-labeled nuclei (*29*, *55*, *56*), as the fewer cases of large-scale automated segmentation of fluorescent membrane-labels were not sufficiently accurate to reliably track the precise shape of each individual cell (*27*, *57*). We show here that, at least in the simple transparent ascidian embryo, a high-resolution morphodynamic atlas of cell geometries can be produced.

This quantitative atlas confirmed the extreme nature of morphological canalization of ascidian development, until the late neurula stage at least, and revealed the quasi-absence of cell migration/intercalation. It also provided the necessary information to build a model to test the hypothesis that local surface-dependent cell inductions collectively exert strong global constraints on embryonic geometries, while changes in the level of expression of inducing ligands were better tolerated.

We propose that this duality between geometric and genetic control of fate specification may be a general phenomenon in animal development. In vertebrates and other embryos developing with high cell number, variants of the “French flag” model of morphogen gradients hypothesize that: “A cell is believed to read its position in a concentration gradient of an extracellular signal factor, and to determine its developmental fate accordingly” (*58*). In this model, individual cell geometries are considered uniform and the main quantitative information is provided by the spatial and temporal shape of the morphogen gradient (Figure 6C). Precisely shaping such gradient, and ensuring their robustness, involves the coordinated recruitment of multiple cellular functions to control the production, degradation, transports or endocytosis of ligands and their receptors (*59*). Even so, the shallowness of such gradient means that inductive cues received by direct neighbors are very similar. The coarse information provided by the gradient is subsequently sharpened over several hours to form clear boundaries through regulatory cross-talk between target transcription factors (*60*) or cell migration. The precise response to morphogen gradients, thus, involves sophisticated layers of regulation, which we propose strongly constrain the architecture of regulatory networks and the evolution of the genome. Modern vertebrate genomes are indeed slowly evolving (*61*).

While neighboring cells may receive very similar signals in a shallow morphogen gradient, they usually share a minority of direct physical neighbors. The surface-dependent read out of ligands we described here thus ensures that the magnitude of the signals they experience differs sufficiently to alleviate the necessity for subsequent transcriptional refinements (Figure 6C). Consistently, fate specification in ascidian occurs in a few minutes within a single cell cycle (*62*), and known gene regulatory networks in ascidians are shallow *(*63*)*. Surface-dependent inductions, in addition to being tolerant to changes in fluctuations in ligand concentrations are therefore likely to involve much fewer layers of regulation, thereby relaxing further constraints on genome evolution.

## Acknowledgements

This work was funded by core support from CNRS to PL, by Inria (core support and IPL Morphogenetics) to CG and GM, by the Geneshape project (ANR-SYSC-018-02) to PL and CG and by the Dig-Em project (ANR-14-CE11-0013-01) to PL, CG and GM. Support was also received from the European Molecular Biology Laboratory (L.H.) and the EMBL Interdisciplinary Postdoc Programme under Marie Curie Actions (U.MF.). L.H. acknowledges support from the Center of Modeling and Simulation in the Biosciences (BIOMS) of the University of Heidelberg. LG was supported by a doctoral contract from the CBS2 doctoral school of the University of Montpellier 2, by the Fondation pour la Recherche Médicale (FRM) (FDT20140931061), and by the Morphoscope2 Equipex project. UMF was supported by the Geneshape project, and by the FRM (SPF20120523969). The Institut de Biologie Computationelle of Montpellier (IBC) supported EF and BL. BL and JL were supported by the Dig-Em project. We thank the members of the Lemaire, Godin and Hufnagel teams for discussion and advice throughout this project, G. Michelin for advice on the segmentation process, and A. McDougall (Villefranche/mer, France) for the generous gift of the PH-GFP expression construct. We are grateful to P. Keller and A. Pavlopoulos (Janelia research labs, USA), and to F. Fagotto (CRBM, Montpellier, France) for critical reading of the manuscript. We also thank the SI2C2 service for IT support and P. Richard and M. Plays for careful animal husbandry. Author contributions are listed in the supplementary materials.

## Supplementary videos

Video S1: 3D rendering of an intensity image of developing Embryo1 after fusion. Vegetal view. Anterior is to the top.

Video S2: Vegetal view of the ASTEC1 segmented embryo. Color code arbitrary. Note the shape of clones produced from individual 64-cell precursors.

Video S3: Vegetal view (left) and side view through a sagittal section (right) of the ASTEC1 segmented and fate colored Phallusia mammillata embryo. Anterior is to the top. Dark red : unrestricted mesoderm, endoderm, and neural progenitors; Dark grey: endoderm; Purple: TLC; Light beige: primary notochord; Light grey: secondary notochord; Dark beige: TVC; Pink: tail muscle; White: mesenchyme; Dark blue: tail epidermis; Bright red: head epidermis; Salmon, light green, light blue, dark green: different regionalized neural plate progenitors; Golden yellow: germ line.

